# Androgen receptor inhibition extends PARP inhibitor activity in prostate cancer models beyond BRCA mutations and defects in homologous recombination repair

**DOI:** 10.1101/2025.01.07.631700

**Authors:** Giuditta Illuzzi, Alessandro Galbiati, Anna D. Staniszewska, Robert Hanson, Chrysiis Michaloglou, Sophie Cooke, Karolina Uznańska, Maja Białecka, Kamil Solarczyk, Harveer Dev, Charles E. Massie, Mark Albertella, Elisabetta Leo, Josep V. Forment, Mark J. O’Connor

## Abstract

Recent phase 3 clinical trial readouts have shown benefit of the combination of poly(ADP-ribose) polymerase inhibitors (PARPi) with androgen receptor (AR) pathway inhibitors (ARPi) in metastatic, castration-resistant prostate cancer (mCRPC). While benefit was particularly evident in patients with tumours harbouring mutations in homologous recombination repair (HRR) genes, improved outcomes were also observed in patients with no such defined alterations in their cancers. Although there is literature linking AR activity with DNA repair pathways, the basis of the interaction between the AR and PARP is unclear. Here, we show that benefit of the combination of ARPi and PARPi in prostate cancer in vitro and in vivo models with no HRR mutations requires ARPi-responsive cells and a PARPi with PARP1-trapping activity, and does not involve an effect of PARPi treatment in modulating the transcriptional role of the AR. Combination benefit is driven by an increase in DNA damage in the form of DNA double-strand breaks and micronuclei formation, which is not due to a direct control of HRR gene transcription by the AR. In addition, we uncover a novel role of PARP1 in modulating AR recruitment to chromatin in the presence of DNA damage. These data shed new light on the interplay between PARP1 and the AR in dealing with genotoxic insults and provide a mechanism of action consistent with the observed clinical benefit of the combination of PARPi and ARPi in patients with prostate cancer.

**Statement of significance:** Combination of androgen receptor pathway inhibitors and PARP inhibitors has shown efficacy in prostate cancer. We provide a mechanistic explanation through increased DNA damage accumulation observed in combination vs single-agent treatments.

## Introduction

The androgen receptor (AR) is the predominant driver of prostate cancer biogenesis and progression. In the metastatic or recurrent state, prostate cancer is normally treated with anti-hormonal therapy or androgen-deprivation therapy. These include first generation AR-pathway inhibitors (ARPi), such as bicalutamide or nilutamide that target AR translocation to the nucleus and prevent downstream signaling. Second-generation ARPi (also known as novel hormonal agents, or NHAs) such as enzalutamide, apalutamide or darolutamide, further improve upon this mechanism, while the ARPi/NHA abiraterone acetate prevents androgen biosynthesis (1).

Poly(ADP-ribose) polymerase (PARP) inhibitors (PARPi) are approved as monotherapy for patients with certain tumour types displaying defects in the DNA homologous recombination repair (HRR) pathway, exemplified by those harbouring mutations in the *BRCA1* or *BRCA2* genes (2). PARPi approved for monotherapy use (olaparib, niraparib, rucaparib and talazoparib) both inhibit PARP enzyme activity and trap PARP proteins at DNA single-strand breaks leading to the accumulation of potentially cytotoxic DNA double-strand breaks (DSB) in HRR deficient backgrounds (3–5). It has been shown that prostate tumours harbour defects in DNA repair genes, most commonly *BRCA2* and *ATM* (6). As such, different clinical trials in metastatic, castration-resistant prostate cancer (mCRPC), have shown the benefit of PARPi monotherapy in patients that progressed on ARPi treatment and whose tumours harbour HRR mutations, leading to their approval in this setting (7,8).

A phase 2 clinical trial investigated the efficacy of the PARPi olaparib in mCRPC in combination with the ARPi abiraterone, showing benefit in unselected mCRPC populations (9). These observations have recently been confirmed in a phase 3 study of olaparib in combination with abiraterone (PROPel study; NCT03732820), and this PARPi-ARPi combination is also supported by trials testing talazoparib in combination with enzalutamide (TALAPRO-2 study, NCT03395197). As perhaps expected, patients with tumours harbouring mutations in *BRCA1*, *BRCA2* or other HRR genes, benefitted most from the PARPi plus ARPi combination, but a benefit in patients harbouring tumours with no such mutations was also reported (10,11). Links between AR activity and the DNA repair machinery have been previously reported (12–16) and are believed to contribute to the observed clinical benefit of combining androgen deprivation therapy with radiation treatment (17). However, the molecular mechanisms behind this interplay are still unclear, particularly in the case of PARPi combinations (18). Here, we sought to gain additional insights by assessing the combination of ARPi plus PARPi in preclinical *in vitro* and *in vivo* prostate cancer models and better understand the key determinants of combination activity.

## Results

### Combination activity between PARPi and ARPi is more pronounced in ARPi-responsive cell lines treated with a PARPi that has effective PARP1-trapping ability

To better understand combination activity between ARPi and PARPi, we performed *in vitro* efficacy studies in a panel of prostate cancer cell lines. We selected cell lines highly sensitive to ARPi treatment (LNCAP, VCAP), others with an intermediate phenotype (C4-2, R1-AD1) and some completely unresponsive to ARPi challenge (R1-D567, LNCAP 95, DU145) (**Supp Fig S1A**). As expected, given the lack of biallelic, pathological mutations in *BRCA1*, *BRCA2*, *ATM* or other HRR genes, none of the cell lines tested displayed high levels of single agent PARPi sensitivity as observed in HRR mutated backgrounds. Specifically, all half maximal inhibitory concentrations (IC_50_) for olaparib in the panel were in the µM range (**Supp Fig S1A**), compared to the low nM IC_50_ values reported for HRR deficient cell lines (19,20).

Combinations were assessed between olaparib and enzalutamide, and synergy scores (21) were derived. Interestingly, scores suggestive of additive combination benefit (synergy score between 0-5) tended to be higher in ARPi-responding cell lines (**Fig 1A**; **Supp Fig S1B-C**). To determine whether this increased combination benefit would also track with PARPi activity, we assessed synergy scores in two of the ARPi-responsive cell lines (LNCAP and VCAP) where we inactivated the *ATM* gene through CRISPR-Cas9 gene editing to increase their sensitivity to PARPi (**Supp Fig S1D**) (20). As shown in **Figs 1B-C**, combination activity was increased in the *ATM* KO backgrounds, compared to their *ATM* WT counterparts (**Supp Fig S1E-F**). Importantly, this increased combination benefit required a PARPi with good PARP-trapping ability (olaparib), as it could not be reproduced by using the weak PARP trapper, veliparib (**Fig 1D**) (19). To determine combination benefit in a different way, we assessed the effect of adding a fixed dose of enzalutamide (response curve is provided in **Supp Fig S1G)**, to a dose-response colony-forming assay experiment with three different PARPi in the C4-2 cell line. In line with our combination matrices data, we observed a decrease of the IC_50_ values of the PARP trappers, olaparib and talazoparib when combined with enzalutamide, while no significant change was observed with the weaker PARP trapper, veliparib (**Fig 1E**). Interestingly, the greatest fold change in IC_50_ of the PARPi used when combined with enzalutamide was with olaparib (**Fig 1F**).

**Figure 1.**
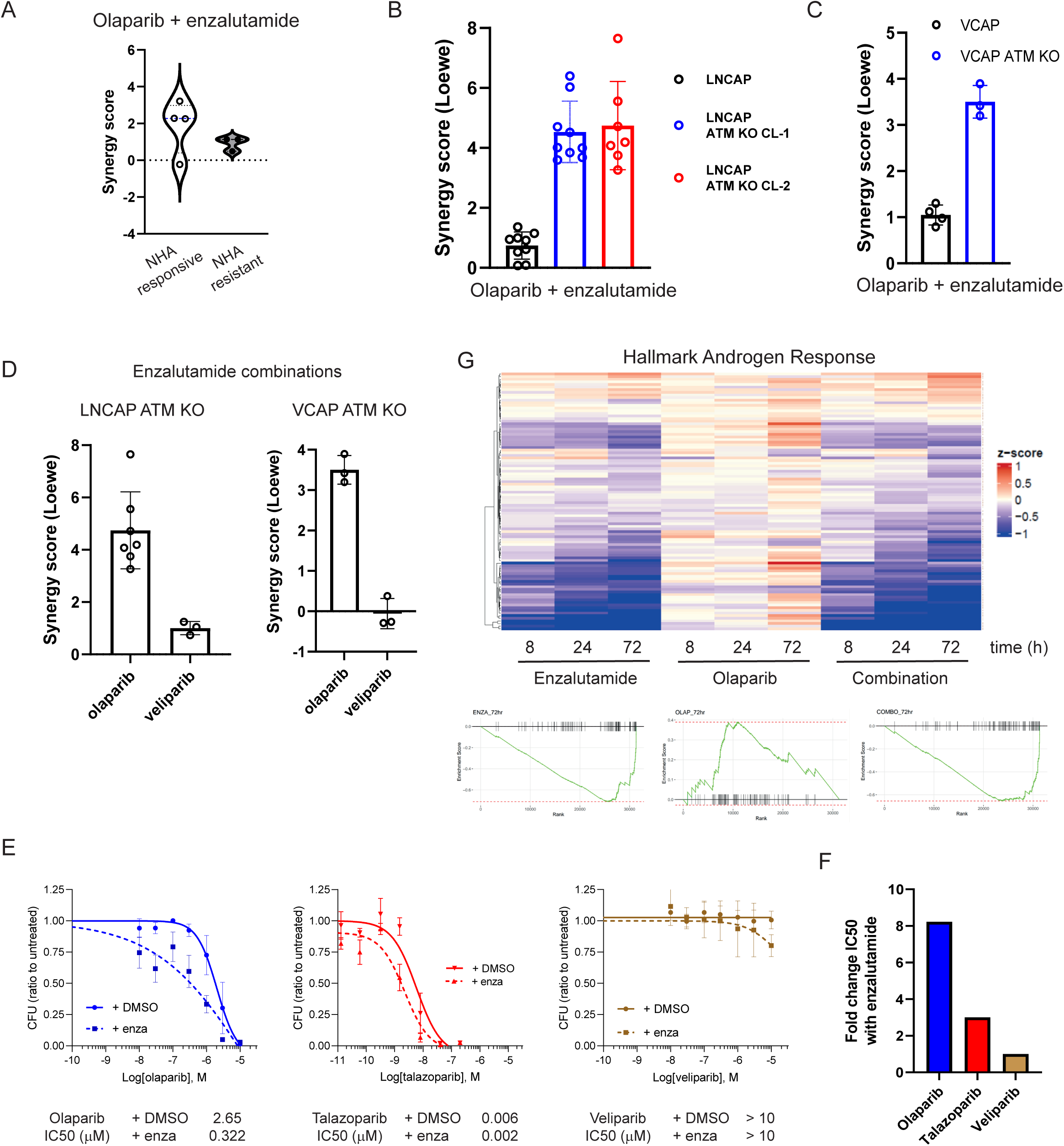
**A)** Combination activity between olaparib and enzalutamide in a panel of prostate cancer cell lines. Each dot represents a cell line (see Supp Fig S1B). **B)** Combination activity between olaparib and enzalutamide in LNCAP and two different isogenic LNCAP *ATM* KO clones. **C)** Combination activity between olaparib and enzalutamide in VCAP and a VCAP *ATM* KO pool. **D)** Combination activity between the PARPi, olaparib or veliparib, and enzalutamide in LNCAP *ATM* KO (left panel) and VCAP *ATM* KO (right panel) backgrounds. **E)** Clonogenic survival assay of C4-2 cells treated with a dose range of different PARPi – olaparib (left panel), talazoparib (middle panel) and veliparib (right panel) – combined with DMSO or a fixed dose of enzalutamide (0.3 μM). IC50 values for the combinations are provided at the bottom of each graph. **F)** IC50 fold change for the PARPi used in panel E) between DMSO and enzalutamide combinations. **G)** RNA-seq differential expression levels in the AR hallmark gene signature in LNCAP treated with enzalutamide, olaparib or their combination (top panel) and leading-edge analysis (bottom panels). In all panels, a minimum of two biological replicates for each condition are presented.

Previous studies have reported that PARPi could synergise with androgen-deprivation therapies by inhibiting a positive transcriptional activation role of PARP1 on the AR pathway (13). For this reason, we were surprised by the lack of combination activity between enzalutamide and veliparib, since this is a potent inhibitor of PARP enzyme activity, in spite of it having weak PARP-trapping ability. Since the reported mechanism of action for the AR-cofactor role of PARP1 does not require PARP1 trapping (13), we decided to explore this previously proposed mechanism further. We assessed the effect of olaparib on the transcriptional profile of LNCAP cells in the presence or absence of ARPi challenge. While enzalutamide treatment had a strong effect at downregulating the expression of AR signature genes, olaparib treatment did not show any modulation of this signature and did not further increase the inhibitory effect of enzalutamide on the expression of this gene set (**Fig 1G**).

Taken together, these results show that combination of PARPi and ARPi *in vitro* is more pronounced in ARPi-responsive cell lines and requires PARPi with PARP1-trapping ability. In addition, combination benefit is higher in genetic backgrounds with increased sensitivity to PARPi, exemplified here by prostate cancer cell lines defective in the *ATM* gene. In addition, our data in **Figs 1E-G** also indicate that PARPi ARPi combination efficacy does not involve any effect of PARPi on AR transcriptional signaling as previously proposed.

### Combination of PARPi with ARPi generates more DNA damage than either single agent

The increased benefit of the combination of PARPi with ARPi was particularly evident when using PARPi with PARP1-trapping activity, pointing towards a DNA damage component driving the combination effect. For this reason, we decided to measure DNA damage by quantifying the accumulation of micronuclei in C4-2 cells treated with enzalutamide, olaparib or their combination. Micronuclei formation is a key measurement of genomic instability and a possible consequence of DNA damage (22). Importantly, combination of enzalutamide with olaparib increased the presence of micronuclei-positive cells compared to either single agent (**Fig 2A**). To confirm that the accumulation of micronuclei could be a consequence of DNA damage, we measured DNA double-strand break (DSB) formation in C4-2 cells treated with enzalutamide, PARPi or their combination using dSTRIDE (23). While enzalutamide treatment did not significantly increase DSB formation, the PARP-trapping PARPi olaparib did, with no significant effect observed with the weaker PARP trapper, veliparib. Strikingly, combination of enzalutamide with olaparib further increased DSB accumulation, an effect that was not recapitulated by the veliparib combination (**Fig 2B**; **Supp Fig S2**). These data correlate with the effect of these combinations in C4-2 cell survival, where enzalutamide increased the efficacy of olaparib but not of veliparib (**Fig 1E**).

**Figure 2.**
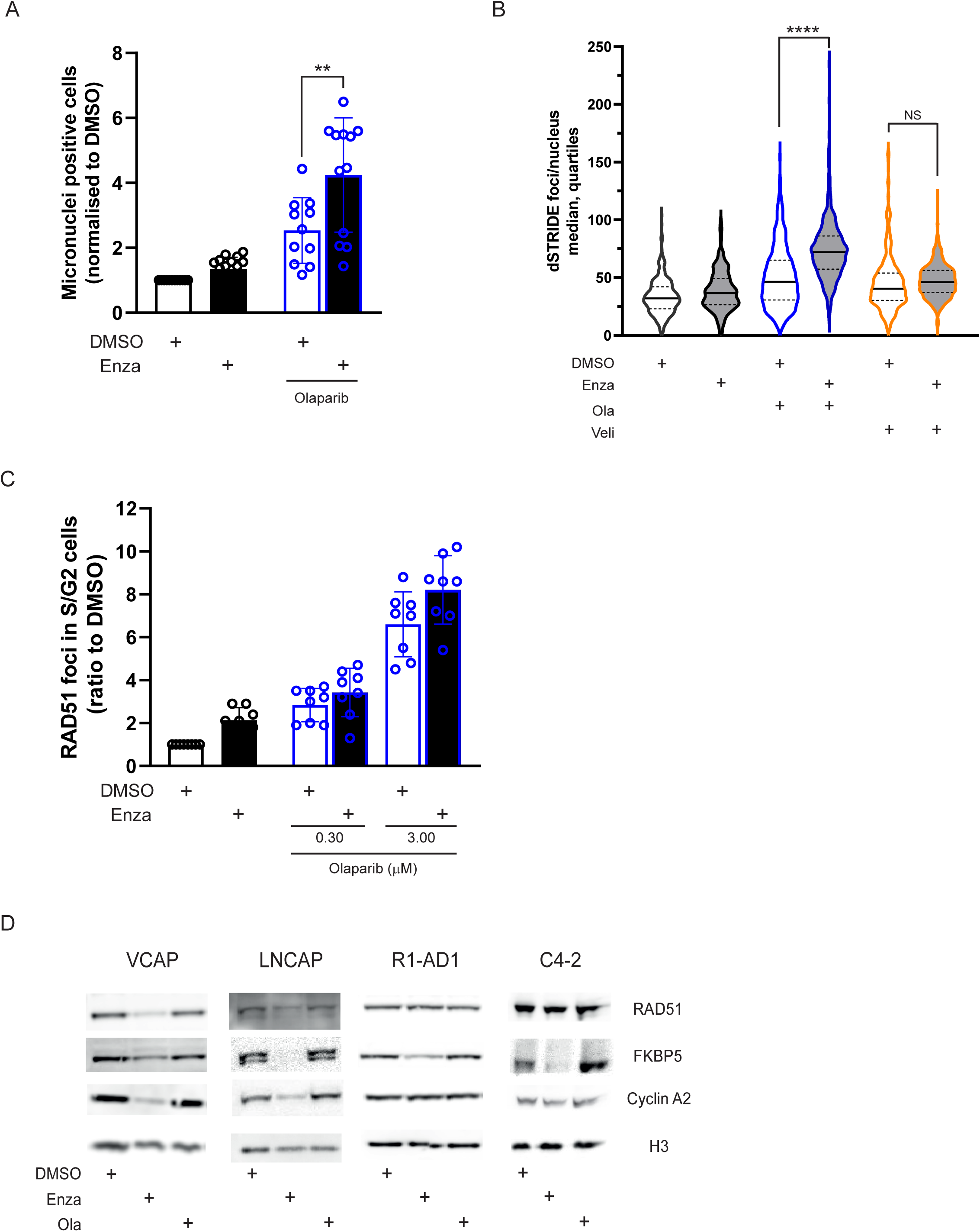
**A)** Quantification of micronuclei in C4-2 cells treated with enzalutamide, olaparib or their combination. Values are represented as ratios compared to DMSO-treated samples. **B)** Quantification of dSTRIDE foci per nucleus in C4-2 cells treated for 72h with 3 µM enzalutamide, PARPi (olaparib or veliparib) 1.8 uM, or their combination. **C)** Quantification of RAD51 foci in C4-2 cells in S/G2 phase of the cell cycle treated with enzalutamide, olaparib or their combination. Values are represented as ratios compared to DMSO-treated samples. **D)** FKBP5, Cyclin A2 and RAD51 protein levels in total cell lysates from VCAP, LNCAP, R1-AD1 or C4-2 cells treated with enzalutamide (3 µM), olaparib (3 µM), their combination or DMSO for 48 h. Histone H3 was used as loading control. In panels A-C, at least n= 3 independent biological replicates for each condition are presented.

Literature reports have suggested that use of ARPi in prostate cancer cell lines could generate a HRR deficient environment, which could explain the ARPi PARPi combination benefit and persistent DNA damage (15,16). To assess this, we measured nuclear focal accumulation of the key HRR recombinase RAD51 (24), as defects in RAD51 foci formation are a *bona fide* readout of HRR defects (20). Olaparib treatment increased RAD51 foci formation in C4-2 cells (**Fig 2C**), as expected in a cell line carrying no defined mutations in HRR genes. Surprisingly, combined treatment of olaparib with enzalutamide did not alter the ability of C4-2 cells to form RAD51 foci, suggesting that no functional defect in HRR was induced by ARPi treatment (**Fig 2C**).

To better understand the discrepancy between our data and previously published observations, we evaluated RAD51 protein expression in VCAP and LNCAP cells, which are very sensitive to enzalutamide treatment, and in R1-AD1 and C4-2 cells, which show a more resistant phenotype (**Supp Fig S1A**). Interestingly, RAD51 protein expression upon enzalutamide challenge was reduced in VCAP and LNCAP cells, while no effect was observed in R1-AD1 and C4-2 cells, despite enzalutamide treatment effectively downregulating protein expression of the AR target gene, *FKBP5*, in all cell lines (**Fig 2D**). We confirmed these observations at the transcriptional level, as enzalutamide treatment caused significant downregulation of the expression of a set of AR-target genes in both LNCAP and R1-AD1 cells (**Supp Fig S3A**), while it only consistently reduced expression of a set of genes involved in the DNA damage response (DDR) in the more sensitive LNCAP cell line (**Supp Fig S3B**).

As expression of a number of DNA repair proteins is downregulated in the G1 phase of the cell cycle, and upregulated during the proliferation phases (S/G2) (25), we investigated the effect of enzalutamide treatment on the cell cycle distribution of these prostate cancer cell lines. Importantly, enzalutamide exposure resulted in very low levels of actively replicating cells in VCAP and LNCAP cells, while much more modest reductions were observed in R1-AD1 and C4-2 cells (**Supp Fig S3C**). This reduced proliferation index was also detected through reduced protein expression of the key S/G2 Cyclin A2 protein exclusively in VCAP and LNCAP cells (**Fig 2D**). Further confirming these observations, enzalutamide treatment significantly reduced expression levels of genes involved in cell cycle regulation only in LNCAP cells (**Supp Fig S3B**). Collectively, these data suggest that rather than direct modulation of HRR genes by ARPi treatment, reduced expression of these genes is an indirect effect of enzalutamide challenge in ARPi-responsive cell lines due to changes in the proliferation status of the cells. This indirect effect of enzalutamide treatment on DDR and cell cycle gene expression, however, did not result in HRR deficiency, as highlighted by the ability of enzalutamide-treated cells to accumulate RAD51 protein at DNA damage sites (**Fig 2C**).

Taken together, these results suggest that the increased efficacy of the ARPi PARPi combination in ARPi-responding cell lines is due to increased DNA damage accumulation in the form of DSBs, driven by the PARP-trapping activity of PARPi.

### The AR is recruited to chromatin in the presence of DNA damage in a PARP1-dependent manner

A direct protein-protein interaction between PARP1 and the AR has been reported (26), so we decided to explore whether such interaction could be modulated in the presence of DNA damage. For that purpose, we performed protein immuno-precipitations of the AR bound to chromatin from C4-2 cells in the presence or absence of olaparib and / or methyl-methane sulphonate (MMS), a DNA alkylating agent that causes DNA damage and strongly induces PARP1 activity. Interestingly, we could observe an interaction between PARP1 and the AR in the chromatin fraction of C4-2 cells, even in unchallenged conditions (**Fig 3A**, lane 2), and this was increased when cells were treated with MMS (**Fig 3A**, compare lane 2 to lane 4) but weakened by the combination with olaparib (**Fig 3A**, compare lane 4 to lane 5). Somewhat surprisingly, the increased interaction between PARP1 and the AR upon MMS treatment was clearly driven by an increase of both PARP1 and the AR in the chromatin-bound fraction (see inputs on lanes 4 and 5 compared to lanes 2 and 3 on **Fig 3A**). This is expected for PARP1, as MMS-induced DNA damage will increase PARP1 trapping on chromatin (19) (note that the reduction on PARP1 levels on **Fig 3A**, lane 5 is driven by the fact that this is a co-immunoprecipitation where the bait is the AR; MMS treatment did induce effective PARP1 trapping, as observed in the input) but it is an unexpected result for the AR. We performed a chromatin-fractionation experiment in LNCAP cells treated with MMS in the presence or absence of enzalutamide, and observed AR recruitment to chromatin after MMS treatment that was reduced by enzalutamide. Importantly, this MMS-induced AR recruitment to chromatin was also affected by PARP1 downregulation using small-interference RNA (siRNA) (**Fig 3B**). These results were confirmed in a chromatin fractionation experiment in C4-2 cells treated with hydrogen peroxide (H_2_O_2_), another potent inducer of PARP1 activity (19). Here, downregulation of PARP1, but to a much lesser extent PARP2, diminished H_2_O_2_-induced AR recruitment to chromatin, which was recapitulated by olaparib treatment (**Fig 3C**). In line with the results using siRNA to inactivate PARP1, PARPi treatment reduced the amount of chromatin-bound AR after MMS treatment regardless of the PARP1-trapping ability of the PARPi used (**Fig 3D**).

**Figure 3.**
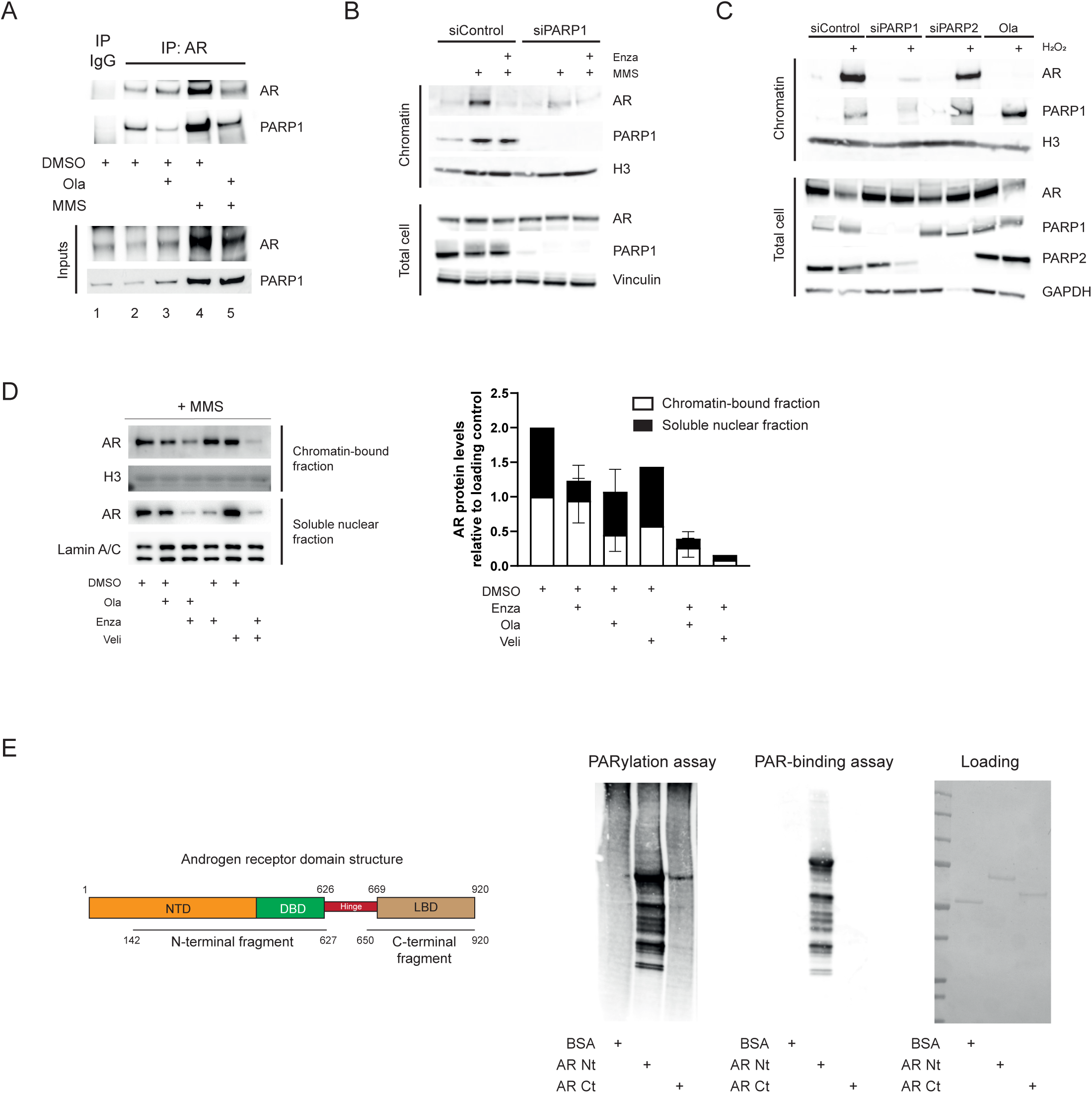
**A)** AR and PARP1 levels in inputs (bottom panels) and AR immune-precipitation (IP) from the chromatin fraction of C4-2 cells treated with MMS (0.01%) for 3 h and pre-treated or not with olaparib 3 µM for 2 h before MMS treatment. **B)** AR and PARP1 enrichment in the chromatin fraction in C4-2 cells treated with MMS (0.01 %) for 3 h and pre-treated with enzalutamide (3 µM) or vehicle control for 16 h. Treatments were added after 48 h incubation with siRNA against PARP1 or a control sequence. **C)** AR and PARP1 enrichment in the chromatin fraction in C4-2 cells pre-treated with olaparib (3 µM) or vehicle control for 2 h before H_2_O_2_ treatment (1 mM) for 30 min. Treatments were added after 72 h of incubation with siRNA against PARP1, PARP2 or a control sequence. **D)** AR nuclear distribution in chromatin-bound or soluble nuclear fractions of LNCAP cells treated with MMS 0.01% for the last 4h after pre-treatment with enzalutamide, olaparib or veliparib (all at 3 µM) as indicated. Left panel: representative western blot images. Right panel: quantification of three independent biological replicates for olaparib and one for veliparib. **E)** PARylation assays with AR N-term or C-term fragments (encompassing domains highlighted on the left figure). Equal amounts of BSA or AR fragments (right panel) were used for a biochemical PARP1-dependent PARylation assay (left panel) or PAR-binding assay (middle panel). NTD = N-terminal domain; DBD = DNA-binding domain; LBD = ligand-binding domain.

Given the physical interaction between PARP1 and the AR, we explored whether the AR could be a direct substrate of PARP1. For that purpose, we purified two different fragments of the AR protein, covering most of the amino acid sequence, and incubated them in the presence of purified PARP1, double-stranded DNA and NAD+, the necessary PARylation cofactor. As shown in **Fig 3E**, PARP1 was able to PARylate the N-terminal fragment of the AR. In addition, we could also observe that the same N-terminal fragment of the AR has the ability to bind PAR chains (**Fig 3E**).

Collectively, these data show that PARP1 and the AR directly interact, and that PARP1 can PARylate the N-terminal fragment of the AR, which can also bind PAR chains. In addition, the AR is recruited to chromatin in the presence of DNA damage in a PARP1-dependent manner.

### Combination activity of PARPi and NHAs in prostate cancer xenograft models

Given the *in vitro* efficacy and DNA damage data generated in ARPi-responsive cell lines, we explored *in vivo* responses to the combination of olaparib with enzalutamide. In LNCAP xenografts, while enzalutamide or olaparib monotherapies had no significant anti-tumour effect, combination therapy resulted in significant tumour growth inhibition (TGI; **Fig 4A**). In C4-2 xenografts, enzalutamide treatment had no significant anti-tumour effect, while olaparib monotherapy resulted in modest TGI. Importantly, combination of enzalutamide with olaparib showed a trend towards enhanced TGI (**Fig 4B**). Taken together, these results show that combination of ARPi with PARPi can result in improved efficacy over monotherapies in prostate cancer xenograft models with no HRR defects.

**Figure 4.**
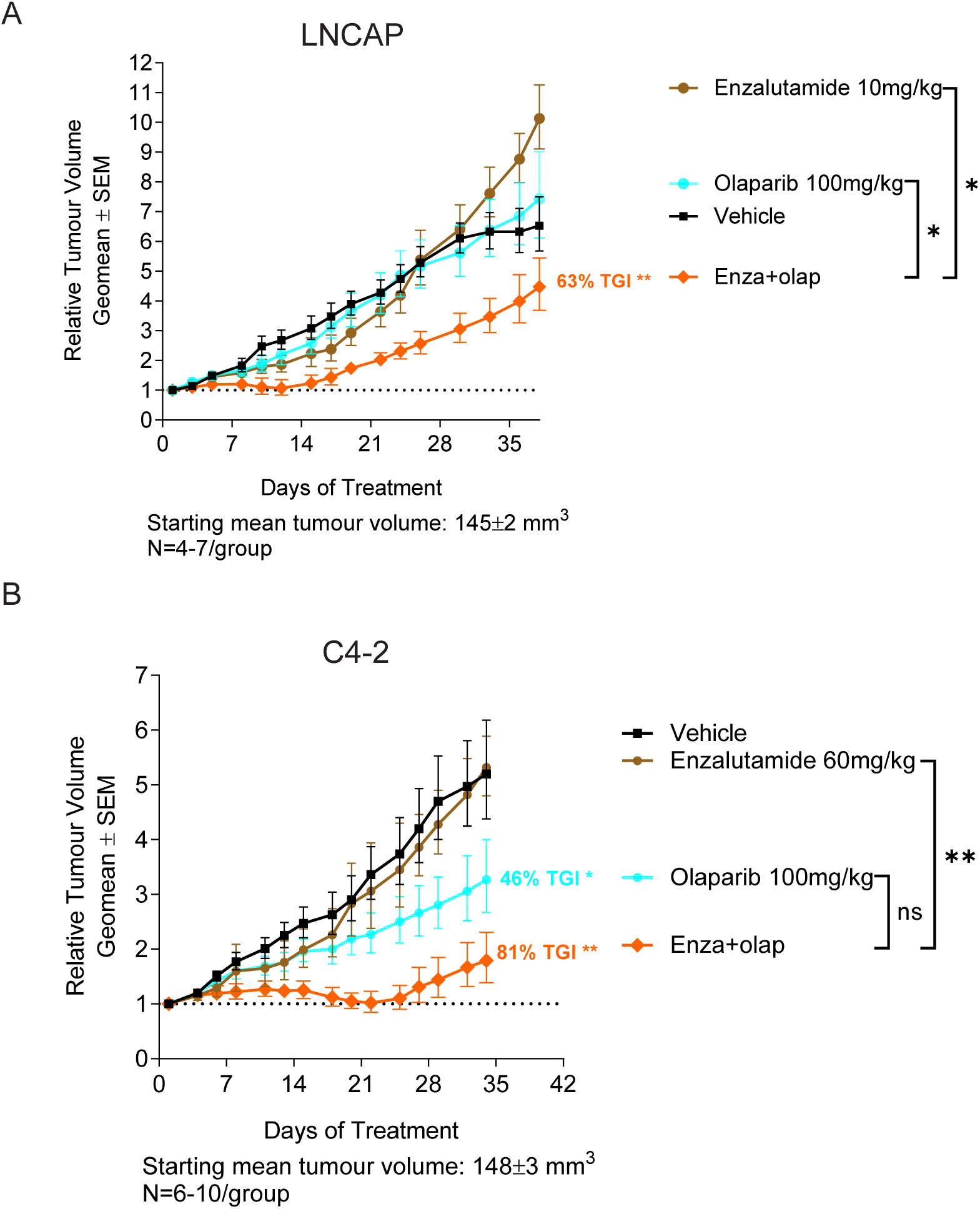
**A)** Anti-tumour efficacy *in vivo* in LNCAP xenograft tumour model treated orally with olaparib (100 mg/kg BID) or enzalutamide (10 mg/kg QD) as monotherapies or in combination. **B)** Anti-tumour efficacy *in vivo* in C4-2 xenograft tumour model treated orally with olaparib (100 mg/kg BID) or enzalutamide (60 mg/kg QD) as monotherapies or in combination. Graphs depict geomean tumor volume ±SEM and percent tumor growth inhibition (TGI). Statistical significance was evaluated compared with the vehicle or combination group using a one-tailed t test (*, P ≤ 0.05; **, P ≤ 0.01; ***, P ≤ 0.001).

## Discussion

PARP inhibitors have demonstrated a clear benefit in HRR-deficient cancers, whether in defined BRCA mutated ovarian, breast, pancreatic or prostate cancers, or in broader HRR gene mutated (HRRm) or HRR-deficient (HRD) cancers defined by genomic scarring criteria. There are still valid questions about the activity of PARPi beyond HRD, particularly as monotherapy. However, the combination of PARPi with ARPi in prostate cancers has been demonstrated in both preclinical non-HRD models, as well as in the clinic in non-BRCA, non-HRR deficient (i.e. HRR proficient) tumours. The mechanism of action of the PARPi ARPi combination behind this activity has been less clear.

In this study, we have shown that the combination of PARPi and ARPi in prostate cancer cell models requires some level of response to ARPi and is further enhanced in the presence of mutations that lead to greater levels of PARPi efficacy. Importantly, we also show that this combination benefit relies on the ability of the PARPi to trap PARP on damaged DNA, and it is not linked to an effect of PARPi treatment on the expression of AR-driven genes. Accordingly, the combination benefit can be explained by an increase in the accumulation of DNA damage in the form of DNA DSBs when PARPi are combined with ARPi. Of note, we report here that this is not due to ARPi treatment generating an HRR deficient environment, and this is in agreement with a recent publication (27). Moreover, we unveil a second component to this combination mechanism of action in the form of the requirement of PARP1 enzymatic activity for the recruitment of the AR to chromatin in the presence of DNA damage.

All these results can therefore be summarized as follows. In untreated conditions (**Fig 5A**), the presence of endogenous DNA damage in prostate cancer cells leads to PARP1 recruitment to chromatin. PARP1 then PARylates itself, the surrounding chromatin and nuclear AR (in abundance in the nucleus in the presence of hormonal signals) to help AR recruitment to chromatin, which in turn will be important to facilitate DNA repair and cell survival. In the presence of ARPi treatment and a PARP-trapping PARPi (**Fig 5B**), PARP1 gets trapped on DNA, increasing DNA damage during replication, and at the same time diminishes AR chromatin binding by inhibiting AR PARylation. ARPi treatment further impairs AR chromatin binding by reducing the total amount of AR in the nucleus. The combination of PARPi and ARPi thus results in an increased amount of DNA damage and leads to enhanced prostate cancer cell death, even in HRR-proficient prostate cancer cells.

**Figure 5.**
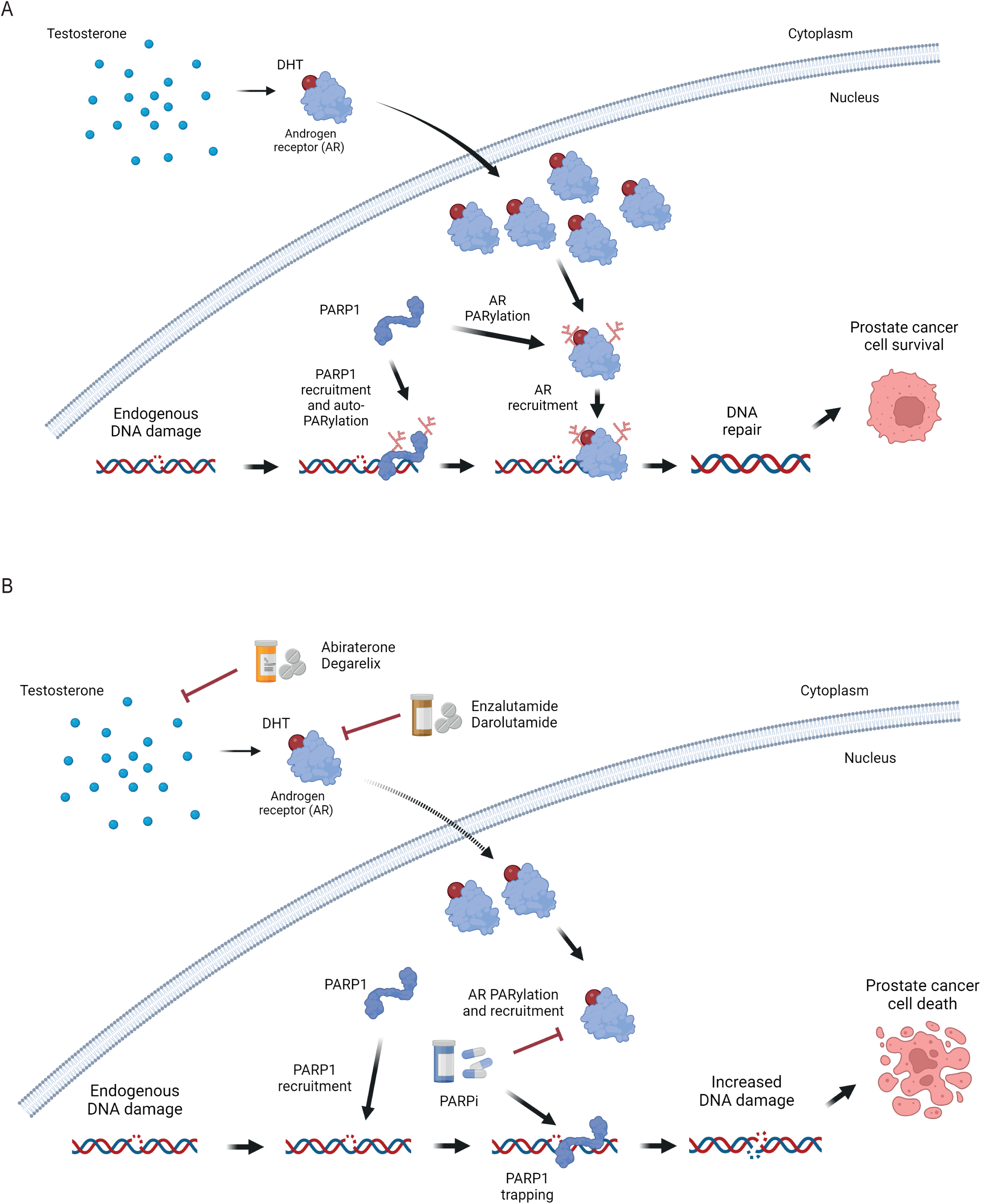
**A)** Model to explain the combined action of PARP1 and the androgen receptor (AR) in facilitating DNA repair in prostate cancer cells. **B)** Model to explain combination efficacy between PARPi and ARPi in prostate cancer treatment. DHT: dihydrotestosterone. Created in BioRender. Forment, J. (2024) https://BioRender.com/q02y075.

This model can help explain the clinical data generated so far with combinations of PARPi and ARPi in prostate cancer and given the importance of the PARP trapping and DNA damage accumulation mechanism for the efficacy of the combination, it explains the lack of benefit of the addition of the weak PARP trapper, veliparib, to the abiraterone regime (28). This mechanism is also the basis of the improved benefit of the combination in HRR mutated backgrounds, including those with mutations in BRCA genes, where PARPi-induced DNA damage will be higher (10,11,29). In non-HRR mutated tumours, this model predicts that being able to maintain effective PARP trapping over the course of the treatment will be needed to generate sufficient DNA damage accumulation to derive benefit in the presence of ARPi therapy. Importantly, dose reductions or dose interruptions of PARPi treatment are reported to result in reduced efficacy and progression on therapy (19). In the PROpel clinical trial, olaparib was maintained at the full monotherapy dose of 300 mg twice a day in most patients (79.9%) and demonstrated benefit in patients with non-HRR mutated tumours when combined with abiraterone (10). In contrast, the niraparib dose when combined with abiraterone in the MAGNITUDE clinical trial, required a dose-reduction of niraparib from its monotherapy dose (from 300 mg once daily to 200 mg once daily), and only demonstrated benefit in patients with HRR mutated tumours (30).

There are limitations to our work that warrant further investigation. For example, we still do not understand the details of how AR recruitment to chromatin in the presence of DNA damage leads to repair, particularly as we did not observe changes in the expression of DNA repair genes by ARPi treatment that cannot be explained by a short-term effect on cell cycle modulation. In addition, the mechanism proposed in Fig 5 does not seem to be applicable to all preclinical prostate cancer models tested, as in some cases we did not detect combination benefit even in ARPi-responsive cell lines. In spite of this, the mechanism for the combination benefit observed between PARPi and ARPi based on increased DNA damage and inhibition of AR-dependent DNA damage repair does draw parallels with the reported clinical benefit of the combination of radiation therapy and androgen deprivation treatment (17). In both cases, the key component is the increased persistence of DNA damage induced by either PARPi or ionizing radiation when the AR function is compromised (12,14). Unfortunately, there are very few prostate cancer preclinical models, in particular those representing relatively common clinical HRR genotypes, and this limits the ability to interrogate these mechanistic interactions in more depth. In order to attempt to overcome these preclinical limitations, a clinical study ASCERTAIN (NCT05938270) has been initiated, where the focus is on collection of pre-and post-treatment biopsies from monotherapy and combinations of the PARP1-selective PARPi, saruparib (AZD5305), and the ARPi darolutamide, with the aim of increasing further understanding of mechanisms of action between PARPi and ARPi. Results of the EvoPAR-Prostate-01 phase 3 clinical trial (NCT06120491), where the combination of saruparib with ARPi is being compared to ARPi monotherapy in metastatic, castration-sensitive prostate cancer patients, will shed further light into the interplay between PARPi and ARPi in both HRR mutated and not mutated settings.

## Materials and methods

### Cell lines and compounds

The following cell lines were originally obtained from ATCC: LNCAP, DU145, PC-3, 22rv1, LAPC-4, VCAP. The isogenic cell lines CWR22Pc-R1-AD1 and CWR22Pc-R1-AD1, were obtained by Dr. Scott Dehm from University of Minnesota (USA). The castrate resistant LNCAP95 (also indicated as LNCAP CR) was obtained by Dr Meeker from John Hopkins University (USA). LNCAP ATM KO clones were as previously described (20). Cell line identification was validated using the CellCheck assay (IDEXX Bioanalytics, Westbrook, ME, USA). All cell lines were validated free of virus and mycoplasma contamination using the MycoSEQ assay (Thermo Fisher Scientific, Waltham, MA, USA) or STAT-Myco assay (IDEXX Bioanalytics). All cell lines were grown in RPMI-1640 growth media (Corning 17-105-CV) supplemented with 10% fetal bovine serum (FBS) or, when indicated, 10% charcoal stripped FBS (ThermoFisher Scientific, 12676029) and 2 mM glutamine. Olaparib, enzalutamide, veliparib, talazoparib, ATM inhibitor (AZD1390) were made by AstraZeneca (Cambridge, UK). Hydrogen peroxide solution was bought by Sigma (cat. No. H1009), methyl methane sulfonate was bought by Sigma (cat. No. 129925). 5-alpha-dihydrotestosterone was bought by Sigma (cat. No. NMID680) and solubilised in ethanol.

### Cell proliferation assay and combination benefit calculation

Cells in 384-well or 96-well plates were dosed using an Echo 555 (LabCyte, San Jose, CA, USA) or using the HP D300e Digital Dispenser (HP Life Science Dispensing), respectively. Cell viability pre-and post-treatment (6-8 days after treatment) was determined using CellTiter-Glo as per manufacturer’s instructions (Promega, Madison, WI, USA; G7570). For the combination experiments data were converted to 0-200% growth inhibition in Genedata Screener as previously described (31) and synergy scores were calculated by comparison to the Loewe model of additivity (21). For three biological repeats, synergy scores and IC50 values were calculated from growth curves averaged from two technical replicates. In the represented plots error bars are mean ± S.E.M for three biological replicates.

### Colony formation assay

Cells were seeded in 24-well plates (1,500 cells/well) and incubated overnight. Drugs were dispensed by an automated digital D300 HP dispenser (Tecan) in titration dilutions, covering 0.1 nM – 10 µM range; each concentration was tested in triplicate and DMSO was used as untreated control. Plates were incubated at 37 °C, 5% CO_2_ for 14 days, to allow colony formation. Cells were then fixed and stained with Blue-G-250 brilliant blue (#B8522-1EA, Sigma, reconstituted in 25% (v/v) methanol and 5% (v/v) acetic acid) for 15min then thoroughly washed with dH_2_O. Plates were scanned with GelCount™ (Oxford OPTRONIX). Colonies were analyzed by total optical density measured with ImageJ software, using a 24-well plate ROI mask. Data analysis was performed by normalization on the vehicle treated of the respective plate set as 1. Data were plotted and IC50s were calculated with GraphPad Prism software.

### RNA-seq

LNCAP cells (5 x 10e5) were seeded in 10 cm dishes and cultured for 3 days before treatment with compounds. Cell lysates were harvested following 8 h, 24 h and 72 h incubation with test compounds, via scraping cells into lysis buffer whilst on ice. RNA was extracted using Qiagen All-Prep kit following manufacturer’s instructions (Qiagen, 2005) and quantified using Qubit RNA BR assay (Life technologies, 2012). 1 μg total RNA from each treatment condition was used as input for mRNA library preparation using the TruSeq RNA Library Prep Kit, following the manufacturer’s protocol (Illumina). The final 36 sample libraries were pooled in groups of 12, forming three equimolar pools each at 10 nM concentration. Sample pools were submitted for paired-end sequencing (2x150bp) on 3 lanes of a HiSeq-4000. Pre-processing of raw read data, alignment and transcript counting was performed using STAR (32) and featureCounts tools (33). Differential expression analysis was performed using the DESeq2 package in R (34). Gene set enrichment analyses (GSEA) were performed on gene lists, ranked by log fold change with respect to DMSO controls, using fgsea package in R (35). Significantly enriched biological pathways were identified through comparison against the Hallmark pathway signatures, from the MSigDB database (36).

### RNA isolation and gene expression analysis by Fluidigm

Cells cultured in 96-well plates were dosed using the HP D300e Digital Dispenser (HP Life Science Dispensing) as indicated. Total RNA was isolated from cells in 96-well plates using the Qiagen FastLane Cell Probe Kit (QIAGEN, 216413), according to the manufacturer’s instructions to a final volume of 40 μL per well, at the indicated time point after treatment.

For Fluidigm analysis cells were snap frozen in RLT buffer and then processed as described in (37). RNA extraction was performed using the RNeasy Plus Mini Kit and RNase-free DNase Kit (Qiagen) on the Qiacube HT instrument (Qiagen) according to manufacturer’s instructions. RNA concentration was measured using the NanoDrop ND8000 (NanoDrop). 50 ng of RNA was used to perform reverse transcription with a Reverse Transcription Master Mix (Fluidigm), and cDNA was then pre-amplified (14 cycles) using a pool of TaqMan primers (Life Technologies) and Preamp Master Mix (Fluidigm). Gene expression analysis was then performed using the 96.96 Fluidigm Dynamic arrays according to the manufacturer’s instructions. Data were collected and analyzed using Fluidigm Real-Time PCR Analysis software. dCt was calculated by subtacting Average Ct of housekeeping genes from each Ct. An average dCt for the control group was calculated and subtracted from dCt to calculate negative ddCt (NegddCt). 2^negative ddCt was used to calculate Fold Change. P values were calculated by performing a student’s t-test on the negative ddCt values. Data were plotted using Spotfire software. Taqman probes used for the Fluidigm analysis are available as on **Table 1**.

**Table 1.**
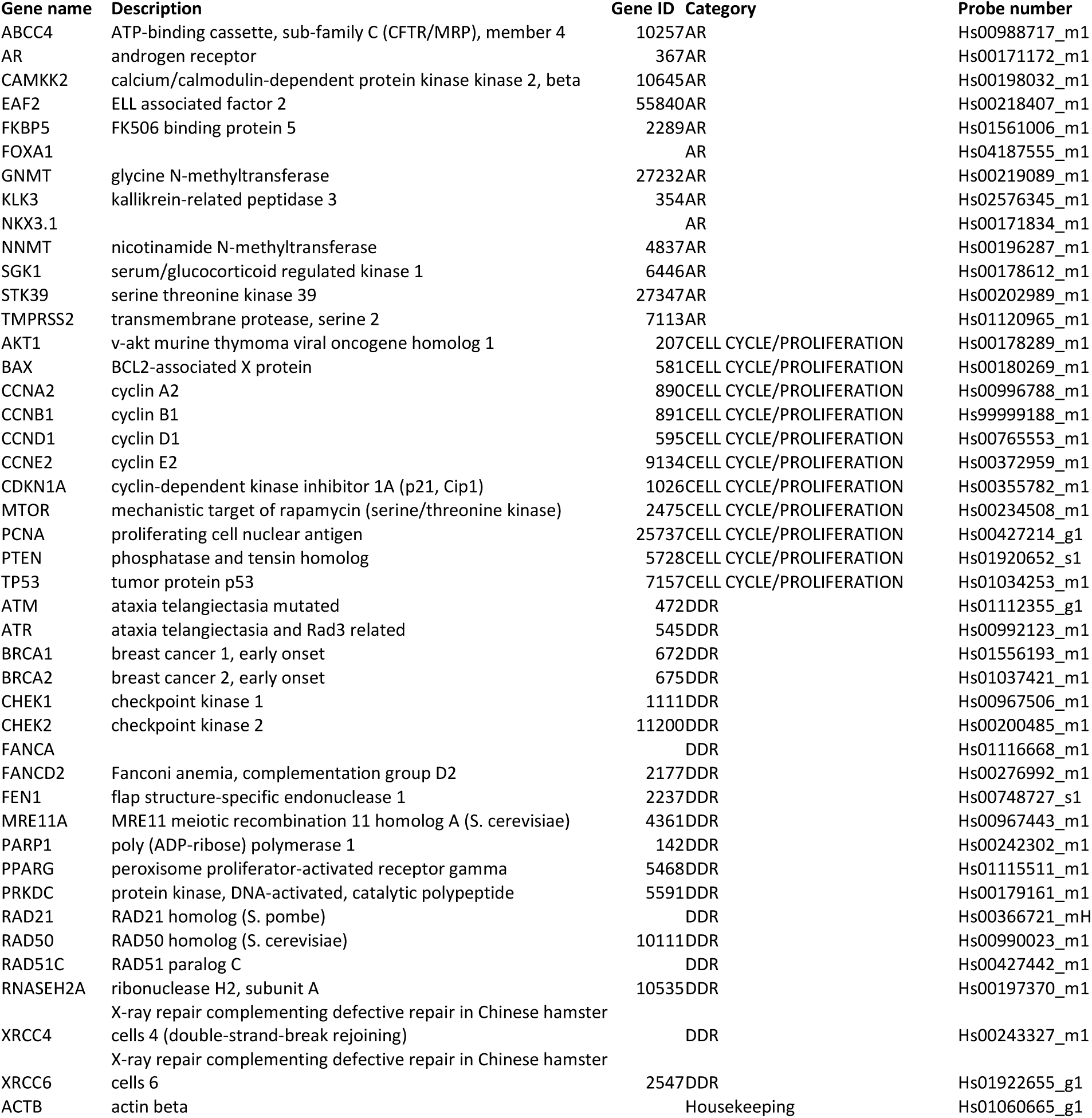
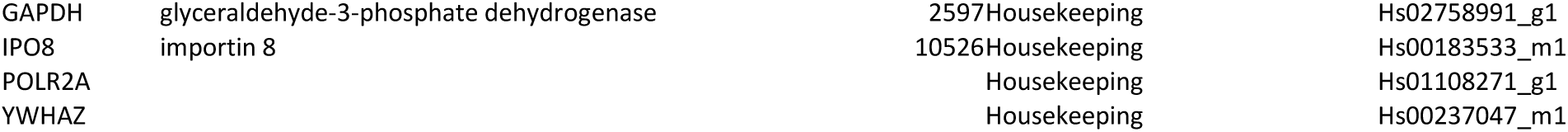
List of Taqman probes used for the Fluidigm studies.

### DNA damage and cell cycle analysis by immunofluorescence

Cells were seeded in imaging-quality 96-well plates (Phenoplates, PerkinElmer #6055300). Compounds were added using a D300 HP dispenser (Tecan) from compound stocks dissolved in DMSO, in duplicate for each condition. Plates were incubated for 48 h. During the last 30 min of incubation, EdU was added to cells at a final concentration of 10 µM, and then processed for immunofluorescence. For immunofluorescence analysis, cells were washed and fixed in 4% paraformaldehyde (PFA) for 15 min at RT and then permeabilized in PBS + 0.2% Triton X-100 for 10 min at RT. Blocking was performed using PBG: 0.5% BSA + 0.2% gelatin from cold water fish skin (Sigma, cat no. G7765) in PBS for 1 h at RT. EdU Click-IT reaction was performed following the manufacturer’s instructions (Thermofisher). Cells were then washed 3 times with PBG; primary antibodies were incubated overnight at 4°C (γH2AX, Merck Millipore, 05-636, diluted 1:5,000. RAD51 antibody, Bioacademia 70-002, rabbit, diluted 1:7,000), followed by secondary antibody (anti-mouse, Alexa Fluor 594 (Thermo, A21099, 1:2,000) and DAPI (ThermoFisher, D1306, diluted 1 µg/ml) for 30 min at RT. Image acquisition and analysis was done with the scanR high content screening platform (Olympus, Evident). 40x objectives were used for image acquisition; 25-36 fields were acquired. For foci analysis “spots detector module” was used. For cell cycle analysis DAPI and EdU signals were used to gate for G1, S, G2 cell cycle phases. For micronuclei analysis an image analysis protocol generated with cellSens(Evident) was used.

### STRIDE assay for detection of DSBs (dSTRIDE)

Direct detection of DSBs was performed at the intoDNA laboratory using the dSTRIDE procedure, as previously described (23). In brief, after overnight fixation of cells in ethanol at -20C, modified nucleotide analogues (BrdU) were incorporated at the sites of DSBs and then detected using appropriate primary antibodies. Next, secondary antibodies with conjugated oligonucleotides and connector oligonucleotides were used and ligated to form a circular DNA template. Finally, rolling circle reaction occurred followed by hybridization of short fluorescently labelled oligonucleotides to the amplicon. STRIDE procedure was followed by DAPI nucleus counterstaining (Thermo Scientific 62248), PCNA IF staining (anti-PCNA antibody Abcam ab29) and CellMask Deep Red cytoplasmic membrane staining (Invitrogen C10046). Fields of view (FOVs) for imaging were chosen randomly over the surface of the sample. The images were then acquired as 3D confocal stacks using Cell Discoverer 7 LSM 900 microscope, Zeiss. Nuclei were recognized based on DAPI staining. 3D masks were created for each recognized nucleus and its projection. Identification of STRIDE foci was based on 3D local maxima of fluorescence inside the nucleus masks. Only the foci localized inside the 3D masks were included in further data analysis.

### Immunoblotting

Cells were lysed in RIPA buffer (Pierce Thermo) supplemented with protease inhibitors (Roche), phosphatase inhibitors (Sigma-Aldrich) and benzonase (Merck, #103773). After 30 min of incubation in ice the lysates were centrifuged at + 4 °C for 15 min and supernatant were kept for sample loading. NuPAGE™ LDS Sample Buffer (ThermoFisher Scientific, NP0008) and NuPAGE™ Sample Reducing Agent (ThermoFisher Scientific, NP0004) were added to the samples. Equal amounts of whole cell lysates were separated on 4-12% Bis-Tris NuPAGE gels and analysed by standard immunoblotting. All immunoblots are representative of experiments that were performed at least twice independently. All antibodies used are described in **Table 2**.

**Table 2.**
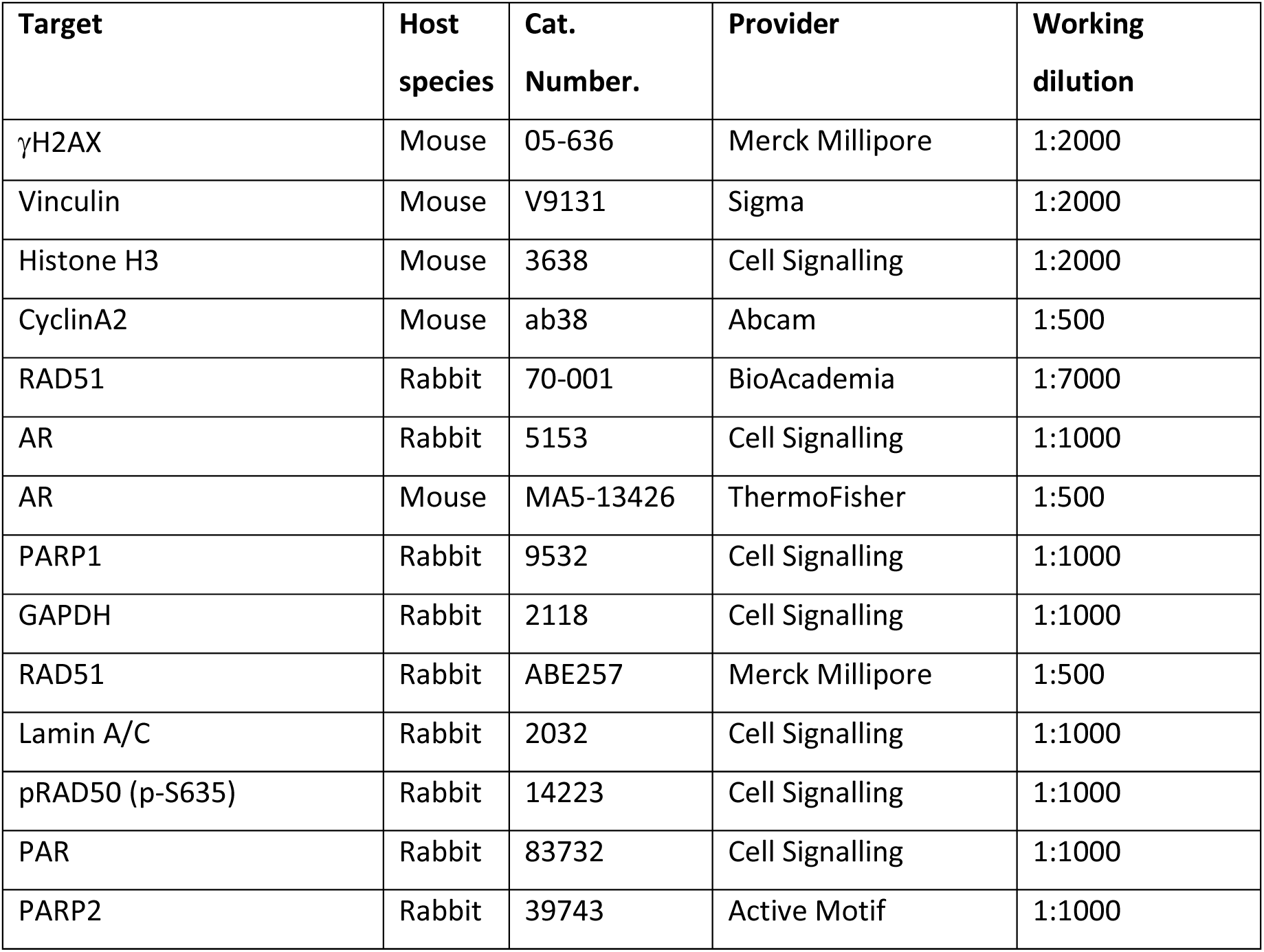
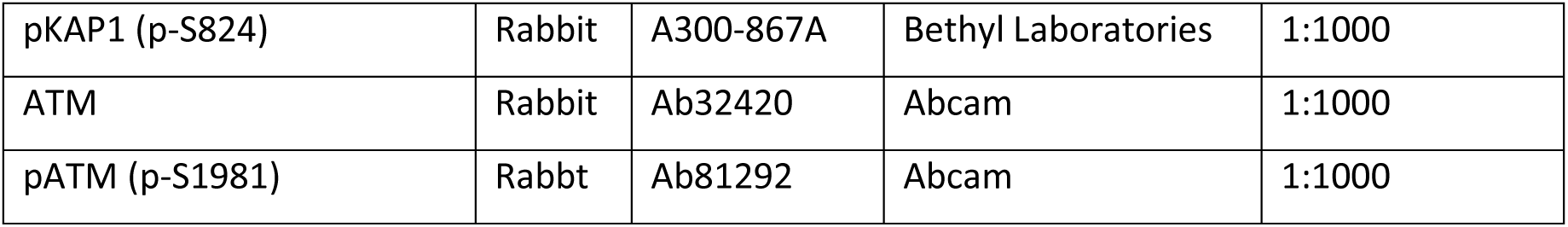
Antibodies for immunoblotting.

### Subcellular fractionation

Cells were plated in 10 cm dishes and treated as indicated. Plates were washed once with PBS and cells were scraped in 1 mL PBS 1x. 20% of the cell suspension was centrifuged and resuspended in RIPA buffer (Pierce Thermo) as described above to obtain the total cell lysate, while the remaining part of the sample was submitted to cellular fractionation using the Subcellular Protein Fractionation Kit for Cultured Cells (ThermoFisher Scientific, 78840) and processed according to manufacturer’s instructions. Equal volumes of each fraction were loaded on SDS gels and processed for immunoblotting as described above. Quantifications were performed with ImageJ image analysis software. For each subcellular fraction AR protein signals were normalised on loading controls (H3 or LaminA/C) and calculated as ratio to DMSO.

### Statistical analysis

Results are shown as mean ± standard error of the mean (s.e.m) or standard deviation (SD) as indicated. P-value was calculated by Student’s two-tailed t-test or with the indicated statistical test, using Graphpad Prism software v8.3.1. Statistical significance is indicated as follows: * p ≤0.05, ** p ≤0.01, *** p ≤0.001, **** p ≤0.0001.

### Gene silencing with siRNA transfection

Cells were seeded in 10 cm plates and transfected with siPARP1 (Horizon Discovery Ltd, Cambridge, UK L-006656-03-0005), siPARP2 (Horizon Discovery Ltd, Cambridge, UK L-010127-02-0005) or non-targeting siCTRL (Horizon Discovery Ltd, Cambridge, UK, D-001810-0X), using the RNAiMax transfection reagents (ThermoFisher Scientific, 13778075), following manufacturer instructions, at a final concentration of 10 nM. Cells were incubated with the siRNA, RNAiMAX, OPTIMEM (ThermoFisher Scientific, 31985070) and the appropriate cell culture medium for 48 h before processing for protein levels and cellular subcellular fractionation assays.

### Immunoprecipitation

The chromatin fraction (50 μL) obtained following the cellular fractionation method described above was diluted in the 500 μL of the IP buffer (20 mM Tris-HCl pH 7.5, 150 mM NaCl, 2 mM MgCl2, 0.5% Triton X-100, 10% glycerol, cOmplete™ Protease Inhibitor Cocktail (Roche). 1:10 of the sample was kept aside as input, while the rest of the sample was incubated over-night at 4 °C, on a rotating wheel, with 2.5 μg of anti-IgG antibody (ThermoFisher Scientific, 31235) or 2.5 μg of anti-AR antibody (CST, 5153). The following day 50 μL per sample of Protein A Dynabeads (10002D) were blocked with PBS + BSA 5mg/mL for 30 min and then washed 3x in the IP buffer. Then, 50 μL of beads were added to each sample and incubated for 3 h at 4 °C, on a rotating wheel. Then, beads were washed 3x in the IP buffer. Finally, the samples were eluted in 50 μL 2x NuPAGE™ LDS Sample Buffer (ThermoFisher Scientific, cat. No. NP0008) + NuPAGE™ Sample Reducing Agent (ThermoFisher Scientific, cat. No. NP0004). Sample Inputs and IP were immunoblotted as described above.

### In-vitro PARylation assay and PAR-binding assay with purified proteins

Purified recombinant PARP1 (6His 6Lys TEV PARP1 1-1014 (Hp155), produced in SF21 insect cell baculovirus vector) and recombinant AR N-terminal (N-His6-TEV-hARV (G142-G627), produced in SF21 insect cell baculovirus vector) were produced by AstraZeneca. AR C-terminal fragment (650–920) was bought from Abcam (ab82124), BSA was used as control (Sigma). For the in-vitro PARP1-dependent PARylation assay, 2.5 μg BSA, 2.5 μg AR N-term or 2.5 μg AR C-term were incubated with the following reaction mix for 30 min at RT: PARP1 (800 ng), dsDNA oligo (5 μg, previously annealed, M3 5’ACTTGATTAGTTACGTAACGTTATGATTGA3’, M4 5’TCAATCATAACGTTACGTAACTAATCAAGT3’), NAD+1mM, Tris HCl pH 7.5 50 mM and water to a total volume of 50 μL per reaction. The reaction was then loaded on a gel and immunoblotted with the primary antibody anti-PAR diluted 1:1000 (CST #83732), as described above. For the PAR binding assay, 2.5 μg of BSA, AR N-terminal and AR C-terminal were mixed with NuPAGE™ LDS Sample Buffer (ThermoFisher Scientific, cat. No. NP0008) + NuPAGE™ Sample Reducing Agent (ThermoFisher Scientific, cat. No. NP0004) and loaded on a on 4-12% Bis-Tris NuPAGE gels. Following gel running and transfer, the membrane was incubated with 100 nM purified PAR polymer (Trevigen, 4336-100-01) for 1 h at RT. Following extensive washing with PBS, the membrane was then processed as a regular western blot and incubated with anti-PAR antibody diluted 1:1000 (CST #83732)

### Xenograft studies

Xenograft studies were performed at Axis Bioservices (Coleraine, UK) in accordance with UK Home Office legislation, the Animal Scientific Procedures Act 1986, Axis Bioservices Animal Welfare and Ethical Review Committee, and the AstraZeneca Global Bioethics policy. Animals were housed in IVC cages (up to 5 per cage) with individual mice identified by tail mark. All animals were allowed free access to a standard certified commercial diet and sanitised water during the study. The holding room was maintained under standard conditions: room temperature at 20-24°C, humidity at 30-70% and a 12h light/dark cycle used. LNCAP or C4-2 cells (10 million with 50% Matrigel) were implanted subcutaneously into the dorsal flank of male non-obese diabetic (NOD) severe combined immunodeficiency (SCID) mice (aged 5-8 weeks, weighing approximately 25-30g) (Axis Bioservices). Tumours were measured (length x width) three times weekly by bilateral Vernier caliper measurements and tumour volume was calculated using the elliptical formula (width × width × length)/2. Animal bodyweight and tumour condition were monitored throughout the study. Mice were randomised into treatment groups when mean tumour volume reached approximately 0.15 cm3. Animals were treated from the day after the randomization. Olaparib (AZD2281, AstraZeneca), and enzalutamide (ChemShuttle) were administered by oral gavage (PO) at 10 mL/kg final dose volume. Olaparib was formulated in 10% DMSO, 30% HB-β-CD. Enzalutamide was formulated in 5% DMSO: 95% Methylcellulose (0.5% w/v) with 0.1% Tween-80. Tumour growth inhibition from start of treatment was assessed by comparison of the mean change in tumour volume of the control and treated groups and represented as percent tumour growth inhibition (TGI, when TV ≥ starting TV). Statistical significance was evaluated using a one-tailed t test. Statistical significance is indicated as follows: * p≤0.05, ** p≤0.01, *** p≤0.001.

### VCAP ATM KO cell line generation

Lentiviral production and transduction was performed in HEK293T cells as described in (20). To establish the VCAP stably Cas9 expressing cell line, VCAP cells were transduced with lentivirus expressing the pKLV2-EF1a-BsdCas9-W (Addgene #67978) vector. The cell pools were then selected with Blasticidine for 3 passages and Cas9 expression was confirmed by western blot. To establish VCAP ATM KO cells, 0.3 x 106 VCAP Cas9 cells were seeded in 6-well plates, and cells were transduced with the lentivirus containing the sgRNA targeting ATM as described in (20). ATM knockout cell pools were analysed by immunoblot and TIDE sequencing to assess the KO efficiency.

