## Supplementary Figures and legends for "Androgen receptor inhibition extends PARP inhibitor activity in prostate cancer models beyond BRCA mutations and defects in homologous recombination repair"

**Supplementary Figure legends**

**Supplementary Figure 1. A)** IC50 values for enzalutamide (left panel) and olaparib (right panel) in the panel of prostate cancer cell lines used in this study. **B)** Combination activity (Synergy score Loewe) between olaparib and enzalutamide for each of the individual cell lines used in this study. Data from a minimum of three independent experiments. **C)** Representative examples of 6x6 combination matrices between olaparib and enzalutamide in each of the prostate cancer cell lines used in this study. In the Fits panel (top), values between 0-100 represent cytostatic effects. The Excess panels (bottom) represent the values used to calculate synergy scores. Values coloured in pink/brown represent positive interactions, while values in blue represent negative interactions. **D)** Characterization of VCAP parental and *ATM* KO pools in terms of ATM-mediated DNA damage signaling by western immunoblotting. Cells were pre-treated or not with ATM inhibitor (ATMi) 2 h before 5 Gy irradiation. Samples were collected 4 h after irradiation. **E)** Representative examples of 6x6 combination matrices between olaparib and enzalutamide in VCAP parental (top panels) and VCAP *ATM* KO pools (bottom panels). **F)** Representative examples of 6x6 combination matrices between olaparib and enzalutamide in LNCAP control (top panels) and LNCAP *ATM* KO clones (bottom panels). **G)** Enzalutamide dose-response in C4-2 cells as measured by clonogenic survival assay.

**Supplementary Figure S2.** Representative images of C4-2 cells treated with DMSO, enzalutamide, olaparib, enzalutamide + olaparib, veliparib or enzalutamide + veliparib and stained with the dSTRIDE protocol. DAPI images are used to detect cellular nuclei for quantification of dSTRIDE foci per nucleus. Distribution of percentages of dSTRIDE foci per nucleus are shown in the histogram plots at the bottom, where each dSTRIDE focus is quantified as a DSB.

**Supplementary Figure S3. A)** Fluidigm analysis of AR-signalling genes in LNCAP (left panel) or R1-AD1 (right panel) cells treated with enzalutamide. Results are shown as mean of n = 3 biological replicates and statistical significance was calculated with a two-sided student's t-test. **B)** Fluidigm analysis of cell cycle- or DNA damage response-related genes in LNCAP (left panel) or R1-AD1 (right panel) cells treated with 3 μM enzalutamide. Results are shown as mean of n = 3 biological replicates and statistical significance was calculated with a two-sided student's t-test. **C)** Cell cycle distribution of LNCAP, R1-AD1, VCAP and C4-2 cell lines treated or not with 3 μM enzalutamide for 48 h.

Supplementary Figure S1

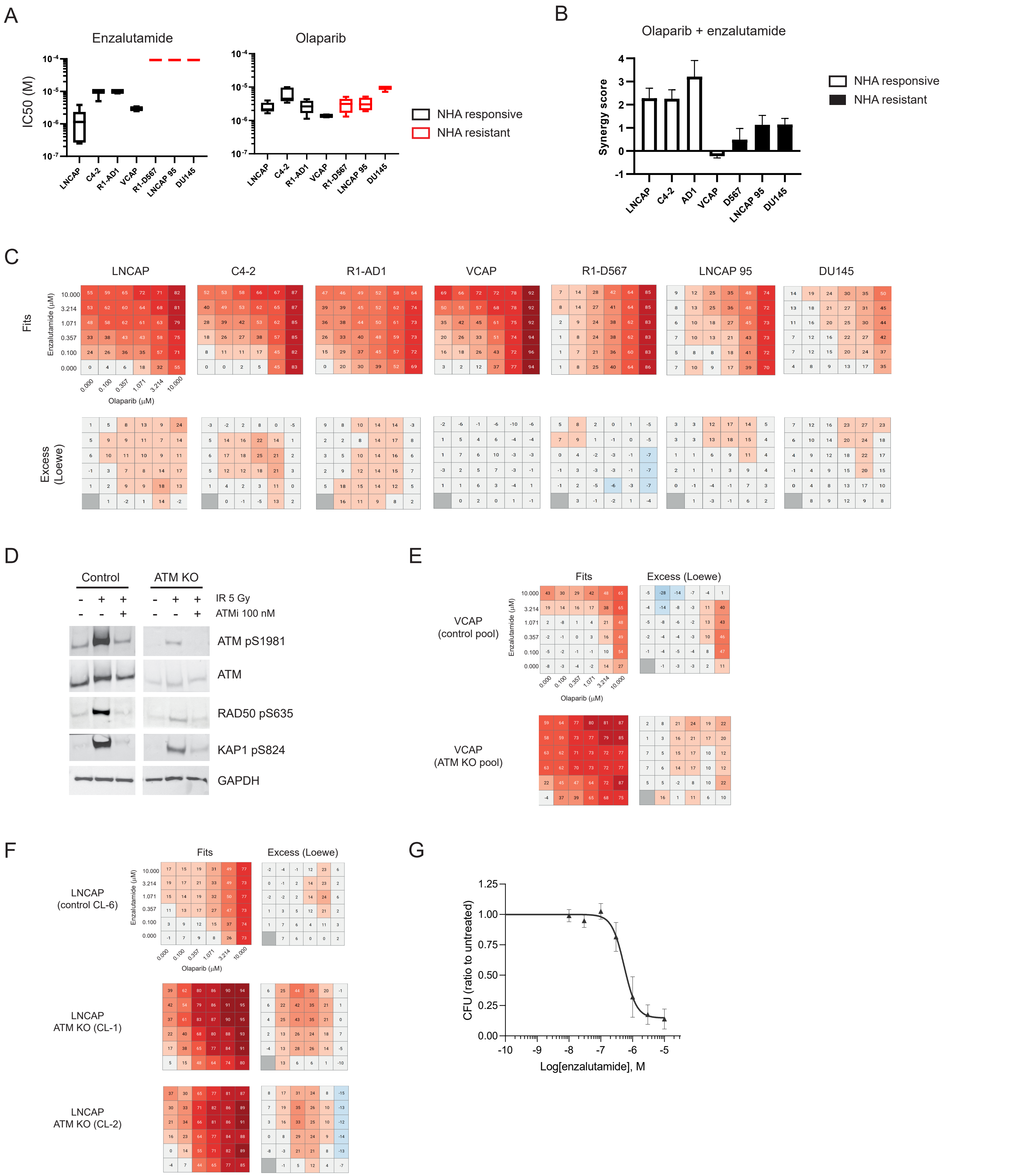

Supplementary Figure S2

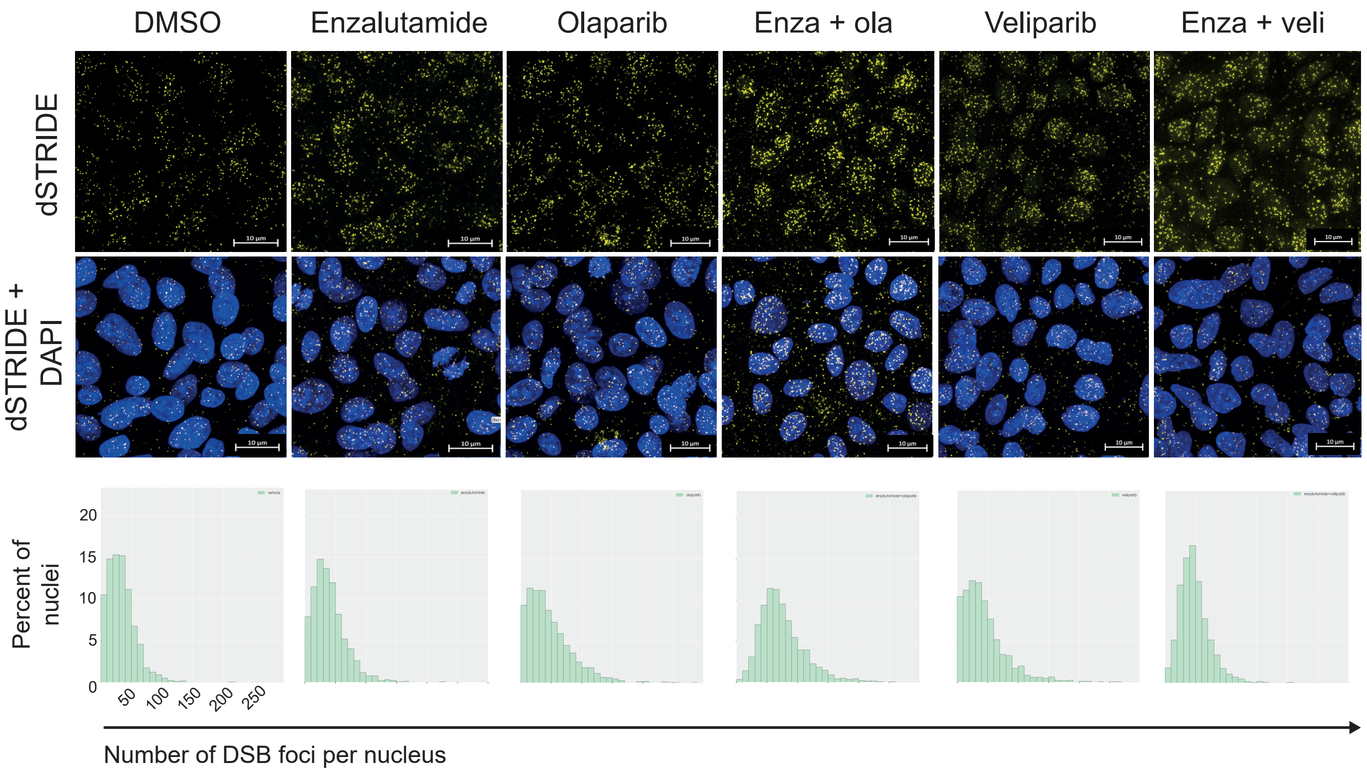

Supplementary Figure S3

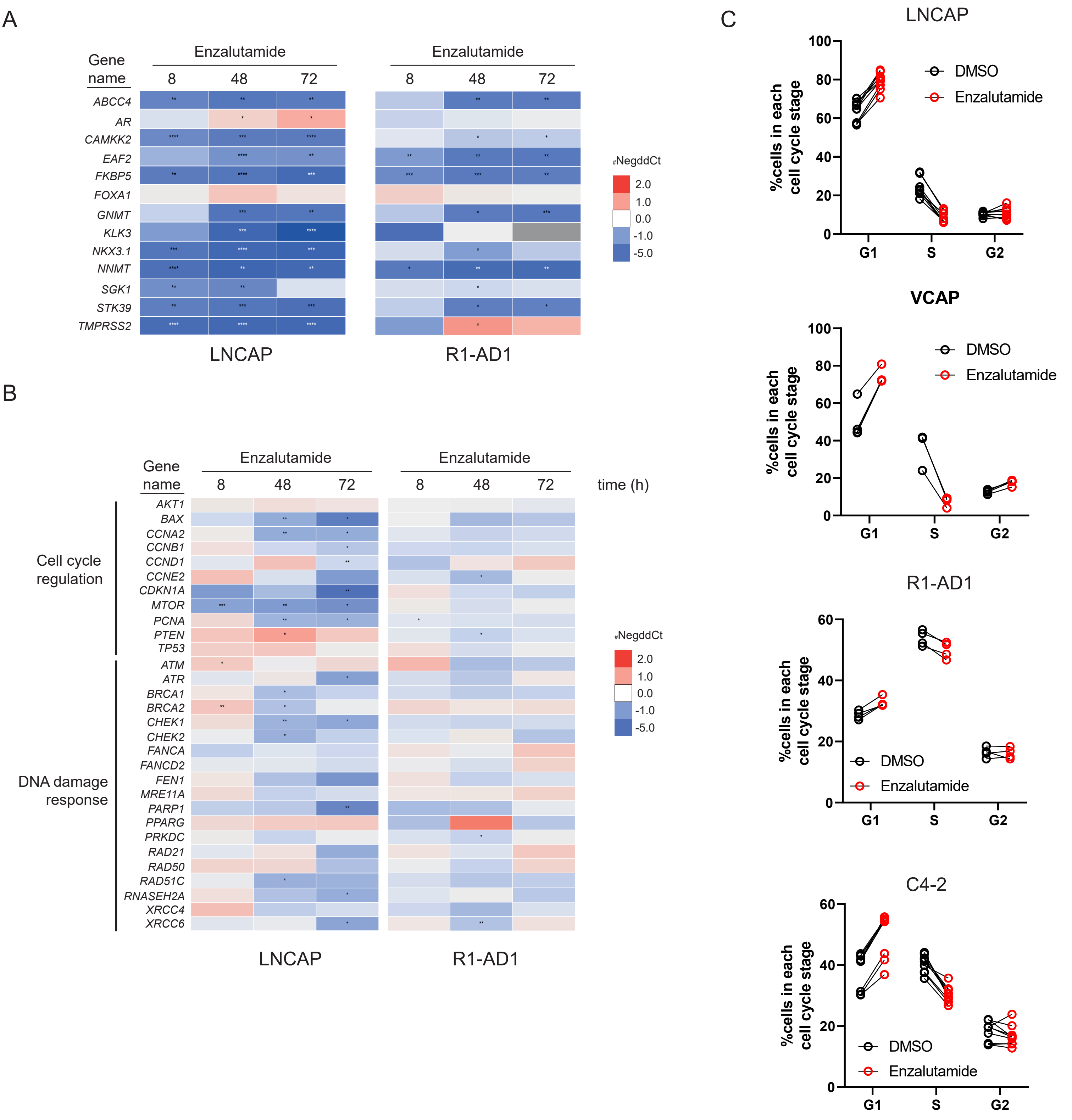
